# Mucosal associated invariant T cells are altered in patients with Hidradenitis Suppurativa and contribute to the inflammatory milieu

**DOI:** 10.1101/2022.01.17.476587

**Authors:** Catriona Gallagher, Julie Mac Mahon, Chloe O’Neill, Féaron C. Cassidy, Hazel Dunbar, Conor De Barra, Caoimhe Cadden, Marta M. Pisarska, Nicole Wood, Joanne C. Masterson, Eoin N. McNamee, Karen English, Donal O’Shea, Anne Marie Tobin, Andrew E. Hogan

## Abstract

Mucosal Associated Invariant T cells are a population of “innate” T cells, which express the invariant T cell receptor (TCR) α chain Vα 7.2-Jα 33 and are capable of robust rapid cytokine secretion, producing a milieu of cytokines including IFN-γ and IL-17. MAIT cells have been reported in multiple human tissues including the gut, periphery and skin. On-going research has highlighted their involvement in numerous inflammatory diseases ranging from rheumatoid arthritis and obesity to psoriasis. Hidradenitis Suppurativa (H.S) is a chronic inflammatory disease of the hair follicles, resulting in painful lesions of apocrine-bearing skin. Several inflammatory cytokines have been implicated in the pathogenesis of H.S including IL-17. The role of MAIT cells in H.S is currently unknown. In this study we show for the first time, that MAIT cells are altered in the peripheral blood of patients with H.S, with reduced frequencies and an IL-17 cytokine bias. We show that CCL20 expression is elevated in lesions of patients with H.S, and MAIT cells can actively traffic towards lesions via CCL20. We show that MAIT cells can accumulate in the lesions from patients with H.S. when compared to adjacent skin, with an IL-17 bias. We show that elevated IL-17, can be linked to the activation of dermal fibroblasts, promoting the expression of chemotactic signals including CCL20 and CXCL1. Finally, we show that targeting the IL-17A transcription factor RORyt robustly reduces IL-17 production by MAIT cells from patients with H.S. Collectively our data details IL-17 producing MAIT cells as a novel player in the pathogenesis of H.S and highlights the potential of RORyt inhibition as a novel therapeutic strategy.

## INTRODUCTION

Hidradenitis suppurativa (HS) is a chronic inflammatory skin condition which affects up to 4% of the population, resulting in the development of nodules and abscesses in apocrine-bearing skin of the axillae, groin, buttocks and the inframammary fold(1, 2). Hidradenitis suppurativa impacts significantly on the patient’s quality of life and emotional well-being(3-5). The pathogenesis of H.S is still not fully understood but emerging evidence has highlighted significant immune dysregulation and inflammation as key drivers of the disease(6). Inflammatory cytokines such as TNF-alpha, IL-1beta and IL-17 have all been identified as possible contributors to H.S(7-10). This is supported by the clinical responsiveness to TNF-alpha inhibitors, corticosteroids and immunosuppressive agents(11-14). However, only a proportion of patients respond to these, thus a deeper understanding of the players and mechanisms underpinning the disease are required. Several immune cell populations have been identified in the lesions of H.S patients, including macrophages, neutrophils, natural killer cells, CD1a-myeloid cells and CD4+ T cells(7, 15-17).

MAIT cells are a population of non-MHC restricted innate T cells that are important in the immune defense against bacterial and viral infections(18). MAIT cells are early responding T cells that are capable of rapidly producing multiple cytokines upon activation such as IFN-gamma, TNF-alpha and IL-17(19). MAIT cells are activated when their invariant TCR recognize bacterial derivatives presented on the MHC like molecule MR1(20). MAIT cells can also be activated in a T cell receptor independent manner, via cytokine stimulation(21). Dysregulated MAIT cell cytokine profiles have been reported in several chronic inflammatory diseases including obesity and arthritis(22-25). Recent studies have identified altered MAIT cells in the skin of psoriasis patients, with an increased frequency of IL-17 producing MAIT cells found in the psoriatic lesion but not healthy skin(26). The authors conclude that MAIT cells are an additional source of IL-17 in psoriatic skin that may be contributing to the disease. MAIT cells have also been implicated in the pathogenesis of dermatitis herpetiformis, a clinical manifestation of gluten hypersensitivity. Frequencies of these cells appear to be significantly increased in this skin disease compared with healthy controls(27). The mechanism is still relatively unknown, however much like psoriasis, IL-17 has been associated with dermatitis herpetiformis pathogenesis(28). To date MAIT cells have not been studied in the context of H.S and were the focus of this study. We show that MAIT cells are altered in the periphery of patients with H.S, with reduced frequencies and elevated production of IL-17. We demonstrate that MAIT cells can traffic via CCL20, the expression of which is elevated in H.S lesions. In the lesions of patients with H.S we observed increased frequencies of IL-17 producing MAIT cells when compared to adjacent skin. Using a cell-based model we show that IL-17 can drive the expression of CCL20 and CXCL1 by dermal fibroblasts, which may support the accumulation of inflammatory T cells and neutrophils in the lesion. Finally, we demonstrate that targeting of RORγt using a small molecule inhibitor limits the production of IL-17 by MAIT cells.

## MATERIALS & METHODS

### Study cohorts

A cohort of 20 adults with hidradenitis suppurativa were enrolled into this study after providing informed consent, along with an aged & sex matched cohort of healthy controls (Table 1). Exclusion criteria included patients with active or recent infections or co-morbid inflammatory conditions outside of H.S. Patients who were on anti-inflammatory medications were also excluded.

### Preparation of peripheral blood mononuclear cells (PBMC) and skin biopsies for flow cytometric analysis

PBMC were isolated from fresh venous blood samples by density centrifugation. Intradermal immune cells (IDCs) were isolated from 6mm punch biopsies of skin. Briefly, biopsies were minced finely with dissection scissors and mixed with 3mL of digestion buffer (0.8mg/mL collagenase type 4 (Worthington) and 0.02mg/mL DNase I (Sigma-Aldrich) in complete RPMI culture media containing 10% FBS). Samples were incubated overnight in 5% CO_2_ before washing through a 100um filter with culture media. CD45 staining was used to identify the immune fraction. MAIT cell staining (1 ×10^6^ PBMC) was performed using specific surface monoclonal antibodies (All Miltenyi Biotec) namely; CD3, CD161, CD8, CCR6, PD-1, CD69 and TCRVα 7.2. Cell populations were acquired using a Attune NXT flow cytometer and analyzed using FlowJo software (Treestar). Results are expressed as a percentage of the parent population as indicated and determined using flow minus-1 (FMO) and unstained controls.

### MAIT cell cytokine production analysis

MAIT cell cytokine production was determined by intracellular flow cytometry or ELISA. Briefly PBMC or IDCs were cultured in presence of a protein transport inhibitor (Biolegend) in media alone (control) or stimulated with cell stim cocktail (Biolegend) for 18 hours at 37°C. After which cells were investigated for intracellular cytokine production (IFNγ and IL-17, both Miltenyi Biotech) by flow cytometry. Purified MAIT cells (50×10^3^) were stimulated with either TCR dynabeads (Gibco) or cell stim cocktail (Biolegend) for 18 hours at 37°C before supernatants where assessed for cytokine by ELISA (R&D systems).

### Tissue and conditioned media analysis

For mRNA experiments, lesions and adjacent 6mm punch biopsies from patients with HS were washed with PBS before isolation of mRNA for analysis by rtPCR. Isolation of mRNA was performed using EZNA Total RNA kit I (Omegabio-tek) according to the manufacturer’s protocol. Synthesis of cDNA was performed using qScript cDNA Synthesis kit (QuantaBio). Real time RT-qPCR was performed using PerfeCTa SYBR Green FastMix Reaction Mix (Green Fastmix, ROX™) (QuantaBio) and KiCqStart primer sets (Sigma). For tissue conditioned media (TCM) experiments, biopsies were washed with PBS before culturing for 18 hours in RPMI. Matched adjacent skin biopsies were used as controls. After 18 hours media was harvested and investigated for expression of CCL20 by ELISA (R&D systems). Interleukin-2 expanded MAIT cells (1×10^6^) from healthy donors were also cultured in the TCM (both control and lesion) for 18 hours with or without TCR Dynabeads (Gibco). Media was harvested and investigated for expression of IL-17 by ELISA (R&D systems).

### MAIT cell trafficking analysis

MAIT cell trafficking was determined using a transwell system. Interleukin-2 expanded MAIT cells (1×10^6^) from healthy donors were plated in the insert (with 3um pore membrane) of a 24-well transwell plate (Corning), with either fresh culture media, culture media containing 20ng/ml of recombinant CCL20 (Biolegend) or conditioned media (either adjacent or lesion). The number of MAIT cells crossing the membrane where enumerated after 4 hours.

### Dermal fibroblast analysis

Human dermal fibroblasts at passage 20 were cultured in complete DMEM (Sigma) supplemented with 1% pen/step (Sigma) and 10% FBS (BioSera). At 80% confluency, cells were treated with recombinant IL-17 (20ng/ml) for 24 hours before harvesting for mRNA extraction and RT-PCR analysis as previously described.

### MAIT cell RORγt analysis

MAIT cell RORγt expression in patients and controls was determined by transcription factor flow cytometry. PBMC, CD45+ skin cells or expanded MAIT cells were activated (TCR dynabeads) for 18 hours before determining RORγt expression using a specific RORγt mAb (BD Biosciences) and transcription factor staining kit (Biolegend). To assess the effect of RORγt inhibition on IL-17 production the specific inhibitor SR1001 (Tocris) was added at a final concentration of 5 μM to purified MAIT cells from either healthy donors or patients with H.S.

### Statistics

Statistical analysis was completed using Graph Pad Prism 6 Software (USA). Data is expressed as SEM. We determined differences between two groups using student t-test or Mann Whitney U test where appropriate. Analysis across 3 or more groups was performed using ANOVA. P values were expressed with significance set at <0.05.

### Study Approval

Ethical approval was granted by the Tallaght University Hospital & St James Hospital Joint Research and Ethics Committee. All patients gave written informed consent prior to partaking in the study.

## RESULTS

### MAIT cells frequencies are reduced in periphery of patients with hidradenitis suppurativa and display an IL-17 bias

We first investigated the frequencies of MAIT cells in the peripheral blood of patients with hidradenitis suppurativa (PWHS), MAIT cells were identified as CD3^+^, TCR Va7.2^+^ and CD161^HI^ after doublet discrimination (Figure 1A). We noted no difference in the percentage of total CD3^+^ T cells between PWHS and controls (Figure 1B). However, we did observe a significant reduction in MAIT cell frequencies in the peripheral blood of PWHS when compared to healthy controls (Figure 1C). We next investigated the activation status of MAIT cells in the peripheral blood PWHS and noted no difference in the expression of CD69, a common T cell activation marker (Figure 1D), but did observe an increase in the expression of the late activation/exhaustion marker PD-1 compared to controls (Figure 1E). Finally, we examined the cytokine production profile and note increased production of IL-17 (Figure 1F) and in parallel decreased IFNγ production (Figure 1G).

**Figure 1:**
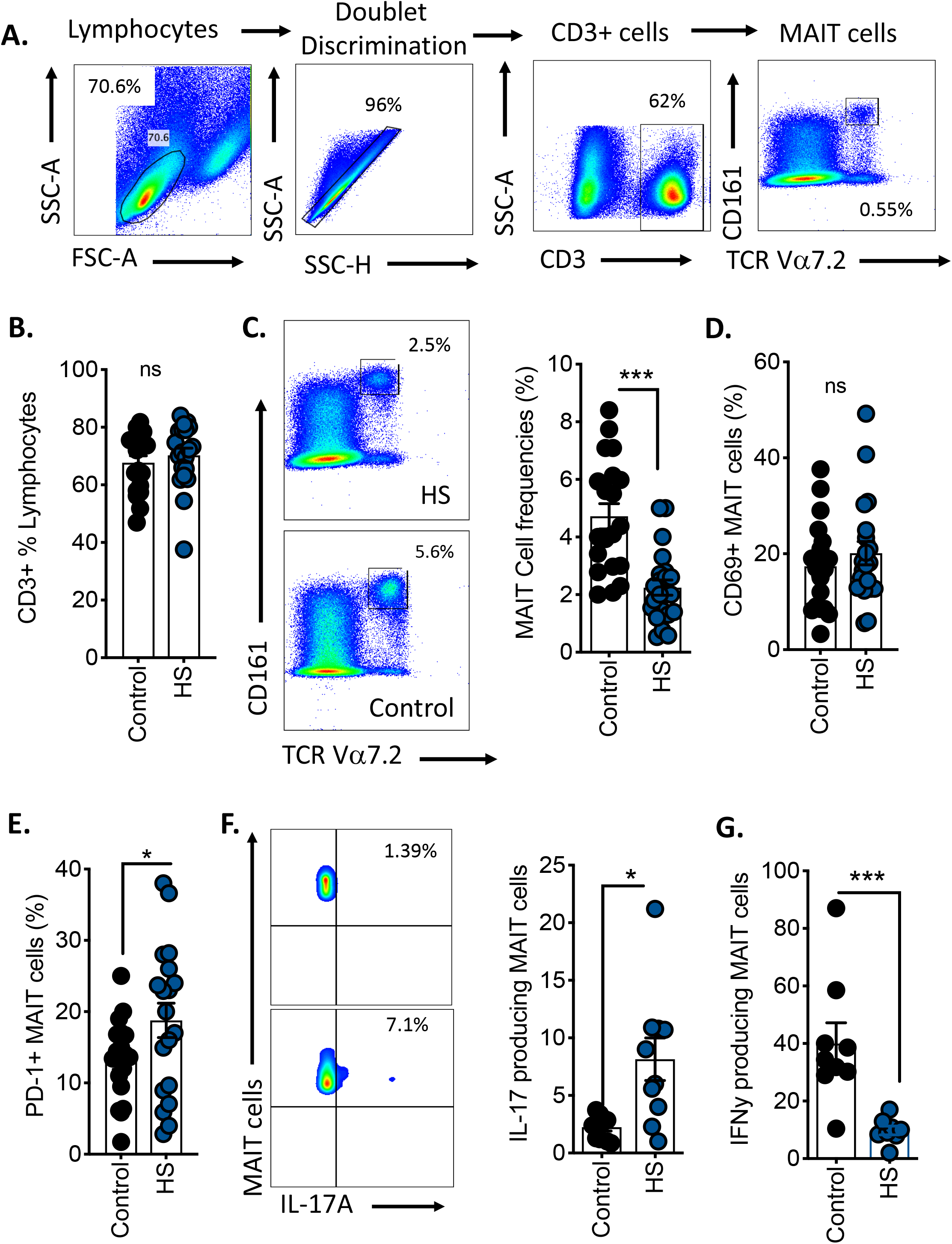
MAIT cells frequencies are reduced in periphery of patients with hidradenitis suppurativa and display an IL-17 bias. **A**. Flow cytometry dot plots outlining the gating strategy for the identification of MAIT cells in peripheral blood. **B**. Scatter plot displaying the frequencies of total T cells (CD3+). **C**. Representative dot plots and scatter plot displaying MAIT cells (CD3+, Va7.2+, CD161+) in peripheral blood of patients with hidradenitis suppurativa or controls (n=10). **D**. Scatter plot displaying the frequencies of MAIT cells expressing CD69. **E**. Scatter plot displaying the frequencies of MAIT cells expressing PD-1. **F**. Representative flow cytometric dot plot and scatter plot displaying the frequencies of MAIT cells (stimulated with cell stim cocktail) producing IL-17 in peripheral blood of patients with hidradenitis suppurativa or controls. **G**. Scatter plot displaying the frequencies of stimulated MAIT cells producing IFNγ in peripheral blood of patients with hidradenitis suppurativa or controls. Data is representative of minimum 5 independent experiments unless otherwise stated. Significant differences are indicated by *(p<0.05), **(p<0.01) and ***(p<0.001).

### MAIT cells can traffic towards the lesions of patients with hidradenitis suppurativa via a CCR6-CCL20 axis

With the observation of reduced MAIT cell frequencies in the peripheral blood of PWHS, we next asked, could MAIT cells be trafficking towards lesions. We first measured the tissue homing chemokine receptor CCR6 expression on MAIT cells from both PWHS and controls and observed no difference in either frequency or intensity (Figure 2A-C). We next measured the levels of CCL20, the chemokine which binds CCR6, in lesions from PWHS and adjacent skin, and observed increased expression of both CCL20 mRNA and protein in lesions when compared to adjacent skin (Figure 2D-E). Finally, using a transwell system, we investigated if MAIT cells could migrate towards either recombinant CCL20 or tissue conditioned media, and demonstrate that MAIT cells actively traffic towards both CCL20 and lesion conditioned media (Figure 2F-G).

**Figure 2:**
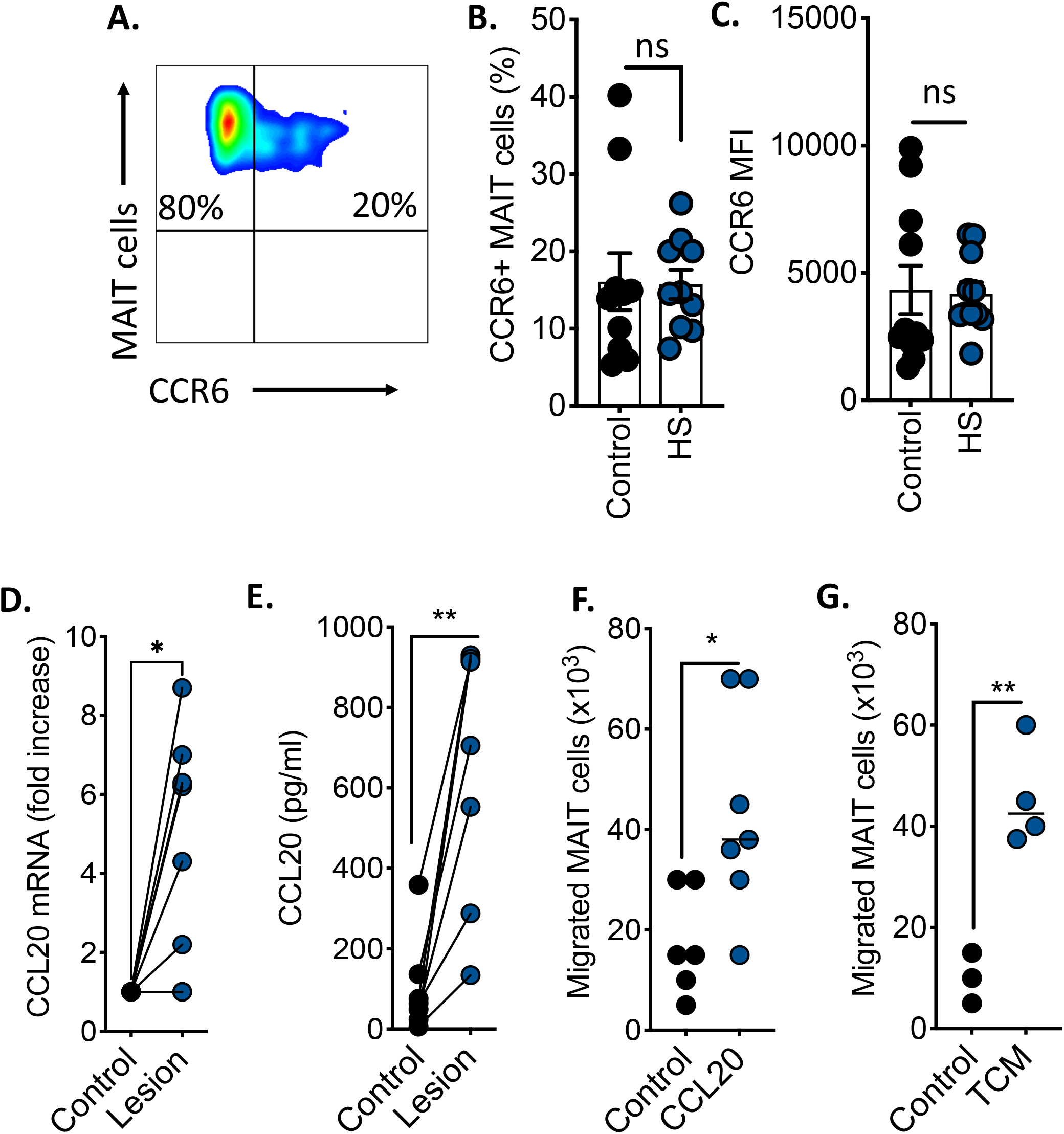
MAIT cells traffic towards CCL20, which is overexpressed in hidradenitis suppurativa lesions. **A**. Representative flow cytometry dot plot showing the expression of CCR6 on MAIT cells from a patient with hidradenitis suppurativa. **B**. Scatter plot displaying frequencies of CCR6% MAIT cells in health controls or patients with hidradenitis suppurativa (n=10). **C**. Scatter plot displaying the mean florescent intensity (MFI) of CCR6 expression on MAIT cells from health controls or patients with hidradenitis suppurativa (n=10). **D**. Line graph showing the CCL20 mRNA expression in HS lesions or matched adjacent skin (controls). **E**. Line graph showing the CCL20 levels in HS lesion or matched adjacent skin conditioned media. F-G. Scatter plots demonstrating the number of MAIT cell migrating across a transwell membrane to either recombinant CCL20 (compared to media alone) or HS lesion conditioned media (compared to adjacent skin conditioned media) Data is representative of minimum 5 independent experiments unless otherwise stated. Significant differences are indicated by ***(p<0.001).

### MAIT cells accumulate in the lesions of patients with hidradenitis suppurativa

Having demonstrated that MAIT cells could migrate towards lesions from PWHS, we next investigated if they were present in lesions from PWHS. The CD3^+^ T cell compartment in skin was identified within the CD45^+^ population, with MAIT cells identified as previously described (Figure 3A). We observed a significant accumulation of MAIT cells in hidradenitis suppurativa lesions compared to matched adjacent skin (Figure 3B-C). Next, we investigated the cytokine profiles of MAIT cells in lesions, demonstrating an increase in MAIT cells producing IL-17 compared to adjacent skin with comparable levels of IFNγ (Figure 3D-F). Finally, we investigated the impact of HS lesion conditioned media on the IL-17 production of healthy MAIT cells and it can drive elevated IL-17 production (Figure 3G).

**Figure 3:**
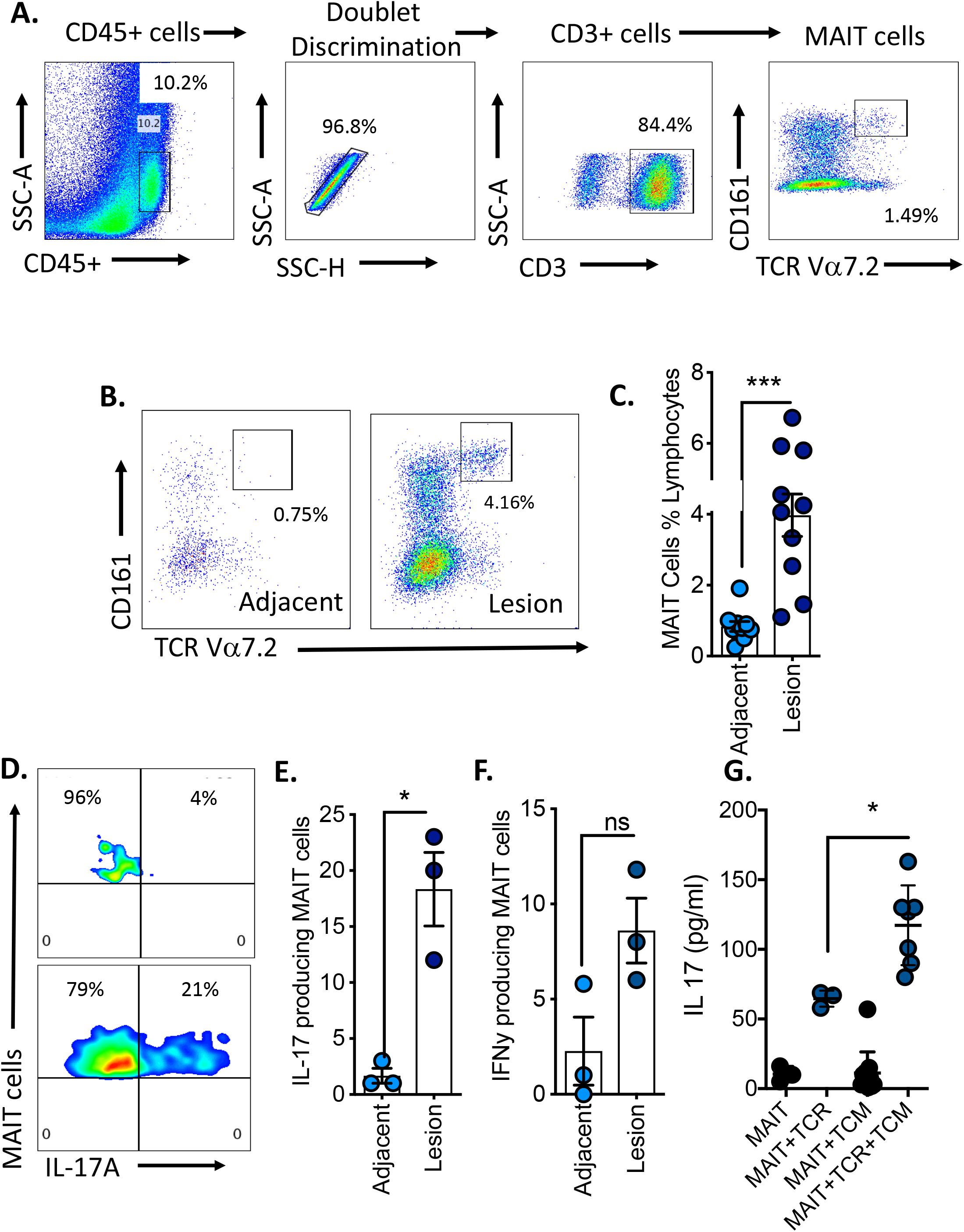
MAIT cells display an IL-17 phenotype in the lesions of patients with hidradenitis suppurativa. **A**. Flow cytometry dot plots outlining the gating strategy for the identification of MAIT cells in skin **B**. Flow Representative flow cytometric dot plot and scatter plot displaying the frequencies of MAIT cells in lesions or adjacent skin of patients with hidradenitis suppurativa. **C**. Scatter plots displaying the frequencies of MAIT cells lesions or adjacent skin of patients with hidradenitis suppurativa. **D-E**. Representative flow cytometric dot plot and scatter plot displaying the frequencies of stimulated MAIT cells producing IL-17 in lesions or adjacent skin of patients with hidradenitis suppurativa (n=3). **F**. Scatter plots displaying the frequencies of stimulated MAIT cells producing IFNγ in lesions or adjacent skin of patients with hidradenitis suppurativa (n=3). **G**. Scatter plot showing the levels of IL-17 produced by expanded MAIT cell lines in the presence of HS lesion conditioned media (denoted TCM), with or without TCR bead stimulation. Data is representative of minimum 5 independent experiments unless otherwise stated. Significant differences are indicated by *(p<0.05), **(p<0.01) and ***(p<0.001).

### Interleukin-17 drives chemokine production by dermal fibroblasts

With the observation of elevated IL-17 in lesions, we investigated the potential impact on dermal fibroblasts using a cell-based model. We showed that IL-17 modulated dermal fibroblast gene expression with reduced Col1A expression but no difference in MMP1 (Figure 4A-B). Interestingly, IL-17 significantly increased the expression of two chemokines, CCL20 and CXCL1 (Figure 4C-D).

**Figure 4:**
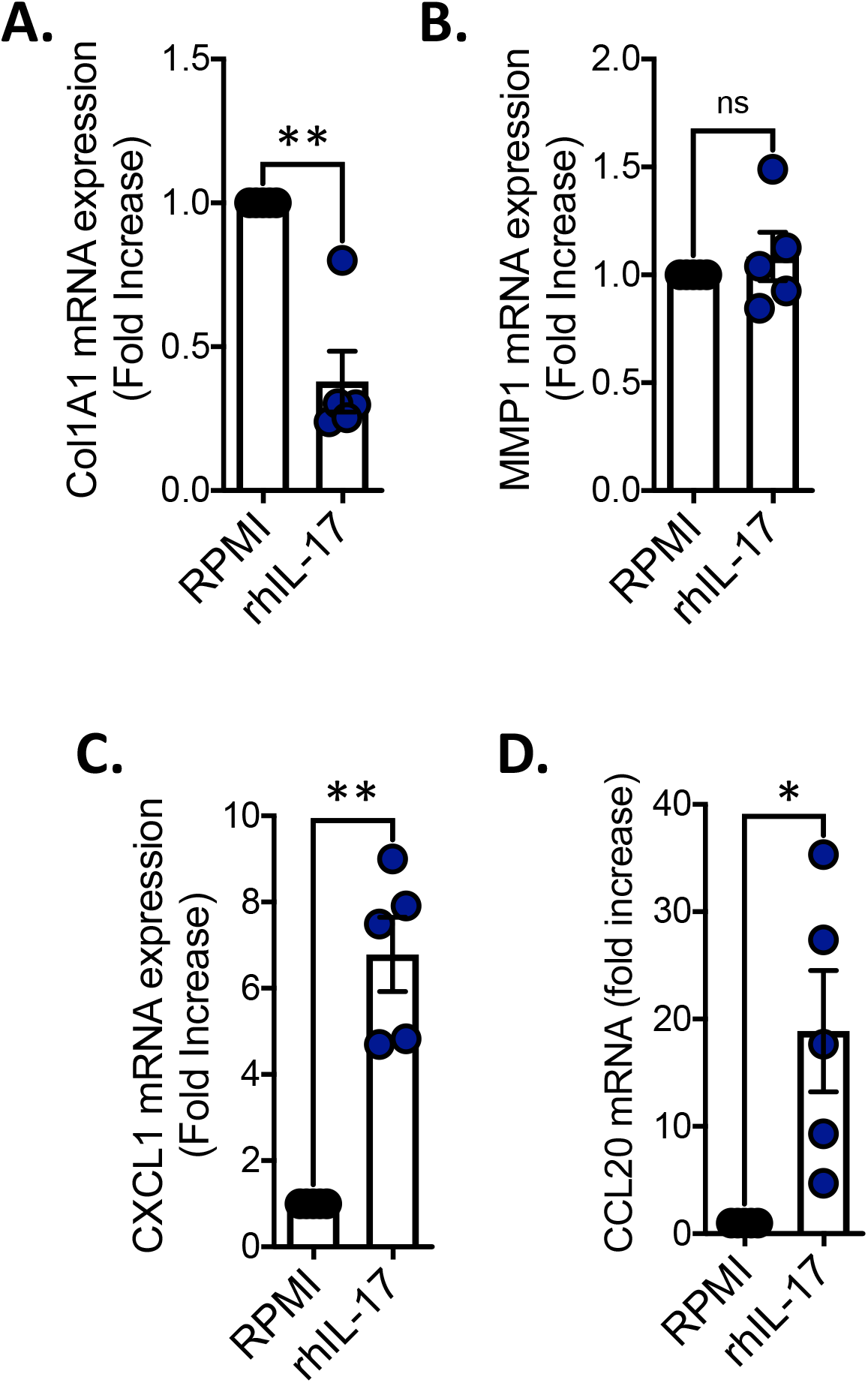
Interleukin-17 drive chemokine production by dermal fibroblasts. **A-D**. Scatter plots displaying the mRNA expression of Col1A, MMP1, CXCL1 or CCL20 in dermal fibroblasts stimulated with IL-17 (20ng/ml) for 24 hours. Data is representative of minimum 2 independent experiments. Significant differences are indicated by *(p<0.05), **(p<0.01) and ***(p<0.001).

### Inhibition of RORyt in MAIT cells from patients with hidradenitis suppurativa results in reduced IL-17 production

IL-17 expression is controlled by the transcription factor RORγt which can be activated via cytokines like IL-23. So, we next investigated the expression of IL-23 mRNA in HS skin and observed elevated levels in lesions when compared to adjacent skin (Figure 5A). We also found that RORyt expression was elevated in MAIT cells from PWHS (Figure 4B-C). We next investigated if targeting RORγt using a specific small molecule inhibitor could reduce IL-17 production by MAIT cells. Firstly, we investigated MAIT cell lines which robustly produced IL-17, and demonstrate that treatment with the RORγt inhibitor SR1001 resulted in reduced IL-17 transcription and secretion (Figure 5D-E). To confirm in the setting of hidradenitis suppurativa we treated samples from patients with hidradenitis suppurativa and demonstrate a robust reduction in IL-17 production (Figure 5F).

**Figure 5.**
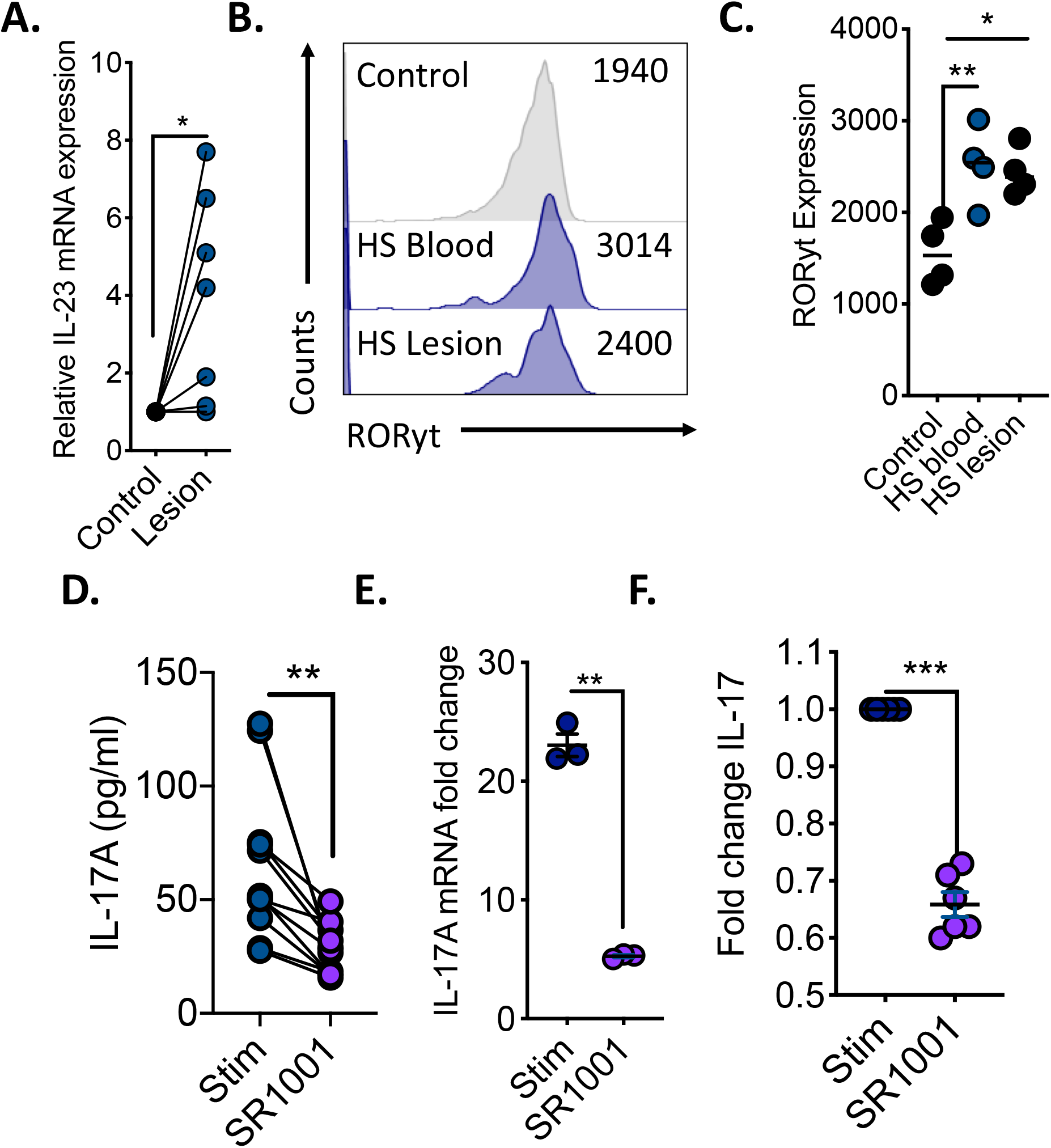
Targeting RORyt in MAIT cells reduces IL-17 in patients with hidradenitis suppurativa. **A**. Scatter plot displaying the mRNA expression of IL-23 in either lesions or adjacent skin (control) from patients with HS. **B-C**. Representative flow cytometric histogram and scatter plot displaying the expression of RORγt in MAIT cells from patients with hidradenitis suppurativa (peripheral blood or lesion) or healthy controls (n=4). **D-E**. Scatter plots displaying the levels of IL-17 (pg/ml) or IL-17 mRNA expression in expanded primary stimulated (TCR beads) MAIT cells with or without SR1001 (5μM) treatment. **D**. Scatter plot displaying the fold change in IL-17 (pg/ml) in MAIT cells from patients with hidradenitis suppurativa treated with SR1001 (5μM) (n=6). Data is representative of minimum 5 independent experiments unless otherwise stated. Significant differences are indicated by *(p<0.05), **(p<0.01) and ***(p<0.001).

## DISCUSSION

MAIT cells are a population of innate T cells which are capable of rapidly producing several cytokines including TNF-alpha, IFN-gamma and IL-17(24). This feature makes MAIT cells potent anti-bacterial and viral effectors but has also implicated them in the pathogenesis of many auto-immune conditions including psoriasis, arthritis and diabetes(26, 29, 30). Hidradenitis suppurativa is a chronic inflammatory skin condition underpinned by a dysregulated immune system, with elevated levels of IL-1, IL-17 and TNF-alpha identified in H.S patients(6). Several immune subsets have been implicated in hidradenitis suppurativa but the contribution of MAIT cells to the inflammatory burden is currently unknown. In this study we show for the first that MAIT cells accumulate in lesions but not adjacent skin from PWHS, this parallels work showing accumulation of MAIT cells in the skin of patients with either psoriasis or dermatitis herpetiformis(26, 27). Similar to inflammatory skin diseases MAIT cells are reported to accumulate in the tissue of several other inflammatory diseases including the joints of ankylosing spondylitis and rheumatoid arthritis patients(31, 32). We show that the chemokine CCL20 is elevated in the lesions of PWHS and that MAIT cells can actively traffic towards CCL20. MAIT cells have been shown to traffic toward CCL20 in other inflammatory conditions such as peritonitis(33). Furthermore, low circulating MAIT cell frequencies have also been associated with elevated CCL20 expression in patients with COVID-19(34). We and many others have highlighted IL-17 producing MAIT cells as potential contributors to pathogenesis of disease(22, 35-37). Here we demonstrate increased expression of the IL-17 producing MAIT cells in both the periphery and lesions of hidradenitis suppurativa patients. We also show that the HS lesion microenvironment could polarize healthy human MAIT cells towards an IL-17 phenotype. IL-17 has been implicated in the pathogenesis of hidradenitis suppurativa (8, 16). We show that IL-17 can alter dermal fibroblasts resulting in elevated CCL20 and CXCL1 expression, both of which have been reported to be increased in PWHS(9, 38). The elevated expression of these cytokines supports the significant immune infiltration reported in PWHS including accumulation of neutrophils and T cells(16, 39, 40). In a recent case study, targeting IL-17 with secukinumab resulted in significant improvements in hidradenitis suppurativa severity(14) highlighting not only a role for IL-17 in disease pathogenesis but a potential therapeutic target. Retinoic acid receptor-related-γt (RORγt) is the key transcription factor in IL-17 producing T cells(41). We assessed the expression of RORγt in MAIT cells from H.S patients and observed increased expression in both the periphery and lesion, compared healthy controls. In 2011, Huh and colleagues were the first to identify small molecular inhibitors of RORγt and demonstrated efficacy in murine models of autoimmune inflammation(42). Since then several studies have investigated the targeted inhibition of RORγt as an alternative to anti-IL-17 biological therapy(43). In spondyloarthritis patients RORγt inhibition selectively targeted IL-17 producing iNKT and γδ-T cells(44). Similarly, Guendisch and colleagues demonstrated positive effects of RORγt inhibition in murine models of arthritis(45). To date the effect of RORγt inhibition on MAIT cells has not been reported, to this end, we assessed the potential impact of RORγt inhibition in primary MAIT cell lines and hidradenitis suppurativa patients, and showed significant inhibition of IL-17. Collectively our data shows for the first time an accumulation of IL-17 producing MAIT cells in hidradenitis suppurativa lesions, and provides supporting data for investigating RORγt small molecule inhibitors in the treatment of patients with hidradenitis suppurativa.

